# The tripartite neural plate border is a common trait of vertebrates and tunicates

**DOI:** 10.64898/2026.07.29.740271

**Authors:** Tasuku Ishida, Yutaka Satou

**Affiliations:** Department of Zoology, Graduate School of Science, Kyoto University, Sakyo, Kyoto, Japan; Ushimado Marine Institute, Okayama University, Setouchi, Okayama, Japan

**Author notes:** Corresponding author: Tasuku Ishida.

## Abstract

The neural crest and neurogenic placodes, which arise from the neural plate border, give rise to morphological characteristics that distinguish vertebrates from invertebrates (Gans & Northcutt, 1983). Recent studies have suggested that embryos of ascidians, a group of tunicates that are the closest invertebrate relatives of vertebrates, possess cells that share an evolutionary origin with vertebrate neural crest cells (Abitua et al., 2012; Fatieieva et al., 2025; Ishida & Satou, 2024; Stolfi et al., 2015; Todorov et al., 2024; Waki et al., 2015) and neurogenic placode cells (Abitua et al., 2015; Ikeda et al., 2013; Liu et al., 2023; Liu & Satou, 2019; Manni et al., 2004; Mazet et al., 2005; Papadogiannis et al., 2022; Wagner & Levine, 2012). To dissect the neural plate border of ascidian embryos at the molecular level, and to gain deeper insights into evolutionary origins of the neural crest and neurogenic placodes, we comprehensively analyzed expression patterns of transcription factor genes at single-cell resolution. We demonstrated that the ascidian neural plate border consists of three domains with distinct gene expression profiles: the anterior, inner lateral, and outer lateral domains. A cross-species comparison of transcriptomes from ascidians and zebrafish suggests that the anterior domain is homologous to zebrafish neurogenic placodes, the inner lateral domain to the neural crest and tail bud and the outer lateral domain to the median fin fold ectoderm. We propose that a tripartite neural plate border was present in the last common ancestor of vertebrates and tunicates, providing a blueprint for evolutionary emergence of vertebrate morphological novelties.

## Introduction

The neural crest and neurogenic placodes contribute to most evolutionary novelties in the vertebrate head, including special sensory organs, cranial ganglia, and jaws (Gans & Northcutt, 1983). Both neural crest and neurogenic placodes develop from the neural plate border, which is located between the neural plate and the non-neural ectoderm. While the neural crest arises from the region abutting the neural plate except for the anterior-most region, the neurogenic placodes originate from the anterior-most neural plate border and the region between the neural crest and the epidermis (Koontz et al., 2023). Although the neural crest and neurogenic placodes have been regarded as vertebrate synapomorphies, embryos of ascidians possess cells that share an evolutionary origin with vertebrate neural crest cells and neurogenic placode cells (Abitua et al., 2012, 2015; Fatieieva et al., 2025; Horie et al., 2018; Ikeda et al., 2013; Ishida & Satou, 2024; Liu et al., 2023; Liu & Satou, 2019; Manni et al., 2004; Mazet et al., 2005; Papadogiannis et al., 2022; Stolfi et al., 2015; Todorov et al., 2024; Wada et al., 1998; Wagner & Levine, 2012; Waki et al., 2015).

In early ascidian embryos, putative developmental counterparts of the vertebrate neural crest and neurogenic placodes emerge from the lateral neural plate border (LNB) or anterior neural plate border (ANB). ANB cells give rise to the adhesive papillae, the oral siphon primordium, and epidermal sensory neurons in swimming larvae (Abitua et al., 2015; Johnson et al., 2024; Nishida, 1987; Nishida & Satoh, 1985) (**Fig. 1a-d**), and are thought to share an evolutionary origin with vertebrate neurogenic placode cells (Abitua et al., 2015; Ikeda et al., 2013; Liu et al., 2023; Liu & Satou, 2019; Manni et al., 2004; Mazet et al., 2005; Papadogiannis et al., 2022; Wagner & Levine, 2012). On the other hand, evolutionary counterparts of the LNB remain extensively debated. The LNB region consists of descendants of b6.5 cells of 32-cell embryos (Ishida & Satou, 2024; Nishida, 1987). At the 64-cell stage, daughter cells of b6.5 (b7.9 and b7.10) are located laterally and adjacent to the neural plate. Among granddaughter cells, b8.17 and b8.19 abut the neural plate, whereas b8.18 and b8.20 are located lateral to b8.17 and b8.19 during the middle and late gastrula stages (**Fig. 1a**). b8.17 and b8.19 cells give rise to ependymal cells of the nerve cord, muscle cells, and tail-tip cells (**Fig. 1d**) (Ishida & Satou, 2024; Nishida, 1987), and likely share an evolutionary origin with vertebrate neural crest cells and neuromesodermal progenitors (NMPs) (Ishida & Satou, 2024). b8.18 and b8.20 cells give rise to epidermal cells, epidermal sensory neurons, and bipolar tail neurons (BTNs) along the dorsal midline (**Fig. 1d**) (Nishida, 1987; Pasini et al., 2006; Stolfi et al., 2015). Evolutionary counterparts of these cells in vertebrate embryos have been debated, with conflicting studies suggesting homology with neural crest cells (Horie et al., 2018; Stolfi et al., 2015; Waki et al., 2015), neurogenic placode cells (Papadogiannis et al., 2022), or the median fin ectoderm (Pasini et al., 2006). Furthermore, pigment-lineage cells (a9.49-lineage cells) that emerged from the neural plate have also been suggested as homologs to neural crest cells (Abitua et al., 2012; Fatieieva et al., 2025; Todorov et al., 2024). In the present study, we comprehensively analyzed spatial gene expression patterns of transcription factor (TF) genes to re-evaluate the homology of neural plate border regions of ascidian embryos with those of vertebrate embryos.

**Figure 1.**
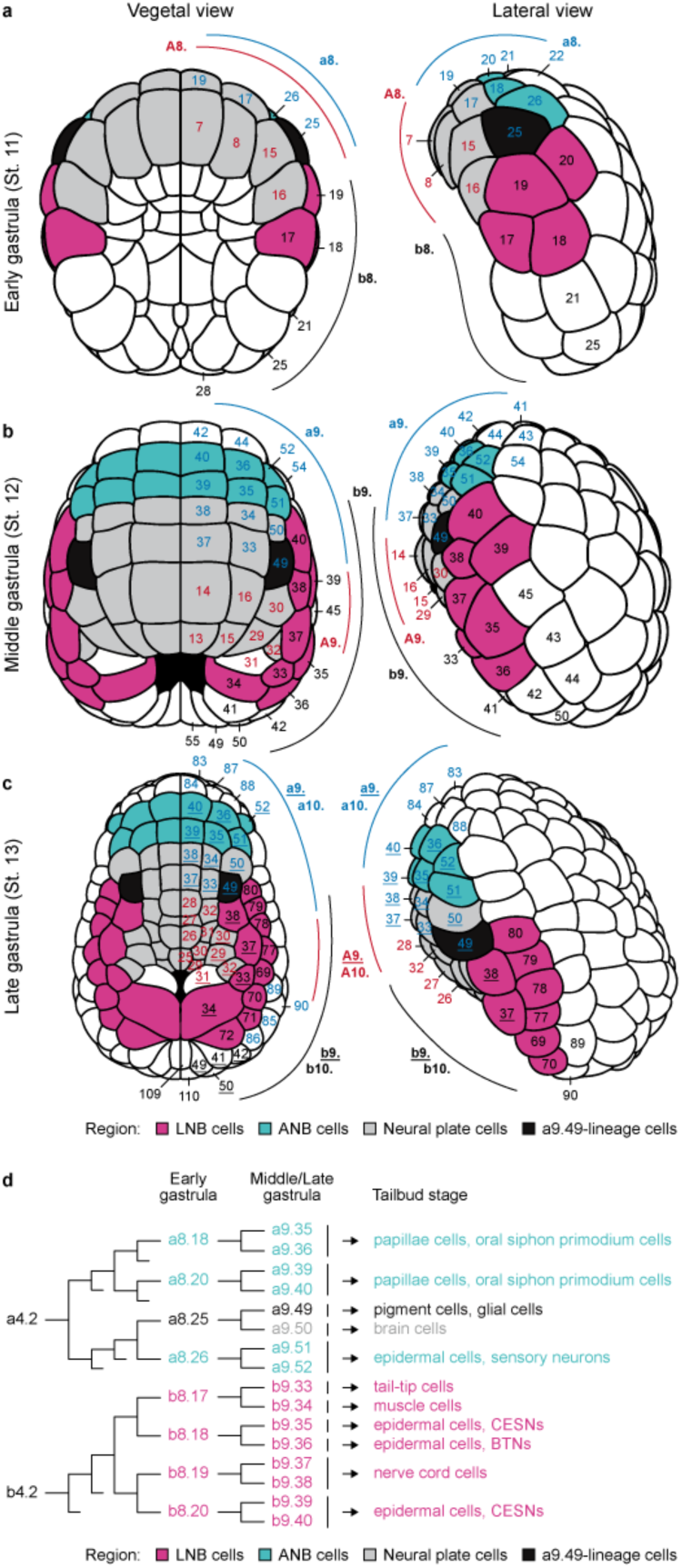
Neural plate border cells in ascidian gastrula embryos. Bilaterally symmetrical ascidian embryos from the early to late gastrula stages. (**a-c**) Schematics of the neural plate border and neural plate cells in ascidian embryos during the early (**a**), middle (**b**), and late (**c**) gastrula stages. Lateral neural plate border (LNB), anterior neural plate border (ANB), neural plate, and pigment-lineage cells are shown in magenta, cyan, grey, and black, respectively. Blastomeres are numbered according to Conklin’s nomenclature, with generations indicated along the margins, e.g., “a9.”. Blastomere numbers are colored as follows: A4.1-lineage (red), a4.2-lineage (blue), and b4.2-lineage (black). In (**c**), underlined and non-underlined numbers denote 9^th^- and 10^th^-generation blastomeres, respectively. (**d**) Cell lineages of LNB, ANB, and neural plate cells. CESNs, caudal epidermal sensory neurons; BTNs, bipolar tail neurons.

## Results

### Comprehensive gene expression profiling of transcription factor genes at single-cell resolution

Specific patterns of TF gene expression, or regulatory states, define cell developmental fates (Davidson et al., 2003). To characterize regulatory states of ascidian neural plate border cells, we comprehensively analyzed spatiotemporal expression patterns of genes encoding TFs using whole-mount *in situ* hybridization in embryos of an ascidian, *Ciona robusta* (formerly *Ciona intestinalis* type A). First, we analyzed publicly available single-cell transcriptome datasets for *Ciona* at the initial and middle gastrula stages (Cao et al., 2019). Fifty-eight TF genes were differentially expressed in the neural plate, neural plate border, and/or epidermis relative to all other cells (**Supplementary Table 1**). Because it is difficult to analyze zygotic expression of genes with abundant maternal expression by *in situ* hybridization, we excluded from the initial list, 11 genes that were expressed abundantly in unfertilized eggs by using publicly available bulk RNA-sequencing datasets from *Ciona* unfertilized eggs (Frese et al., 2024) (**Supplementary Table 1**). In addition, because 17 TF genes that were not included in the list are also expressed in these regions (Abitua et al., 2012; Coulcher et al., 2020; Ikeda et al., 2013; Imai et al., 2004, 2006; Liu & Satou, 2019; Papadogiannis et al., 2022; Wagner & Levine, 2012; Waki et al., 2015), we included these genes as well. We also added 23 *Ciona* orthologs of TF genes that are expressed in the neural plates and neural plate border of vertebrate embryos (Betancur et al., 2010; Gómez-Skarmeta et al., 2003; Simões-Costa & Bronner, 2015; Thawani & Groves, 2020; B. Thisse et al., 2004). Consequently, the final list included 73 TF genes as candidates for patterning the ectoderm in *Ciona* embryos (**Supplementary Table 1**).

We then analyzed spatiotemporal expression patterns of these 73 TF genes by whole-mount fluorescence *in situ* hybridization during the early, middle, and late gastrula stages, and detected clear zygotic signals for 59 of the 73 (**Figs. 2,3; Supplementary Figs. 1,2**; **Supplementary Table 2**). Figure 2 shows expression patterns of *Myt1*, *Otx*, and *Ebf*. During the early gastrula stage, *Myt1* was expressed in the A7.6 cell pair (endomesodermal cells) and a subset of neural plate cells (**Fig. 2a,b**). By the middle and late gastrula stages, *Myt1* was expressed in a subset of neural plate cells, epidermal cells, and the b9.37 cell pair (presumptive nerve cord cells) (**Fig. 2a,b**). *Otx* was initially expressed in LNB cells, ANB cells, a subset of neural plate cells and endomesodermal cells during the early gastrula stage, as previously reported (Hudson & Lemaire, 2001). Then, it was expressed in ANB, neural plate, and epidermal cells at the middle and late gastrula stages. While *Ebf* expression was not observed during the early gastrula stage, it was observed in a subset of neural plate cells during the middle to late gastrula stages. Expression patterns of the 73 TF genes are summarized in Figure 3 for the middle gastrula stage, and in Supplementary Figure 2 for the early and late gastrula stages.

**Figure 2.**
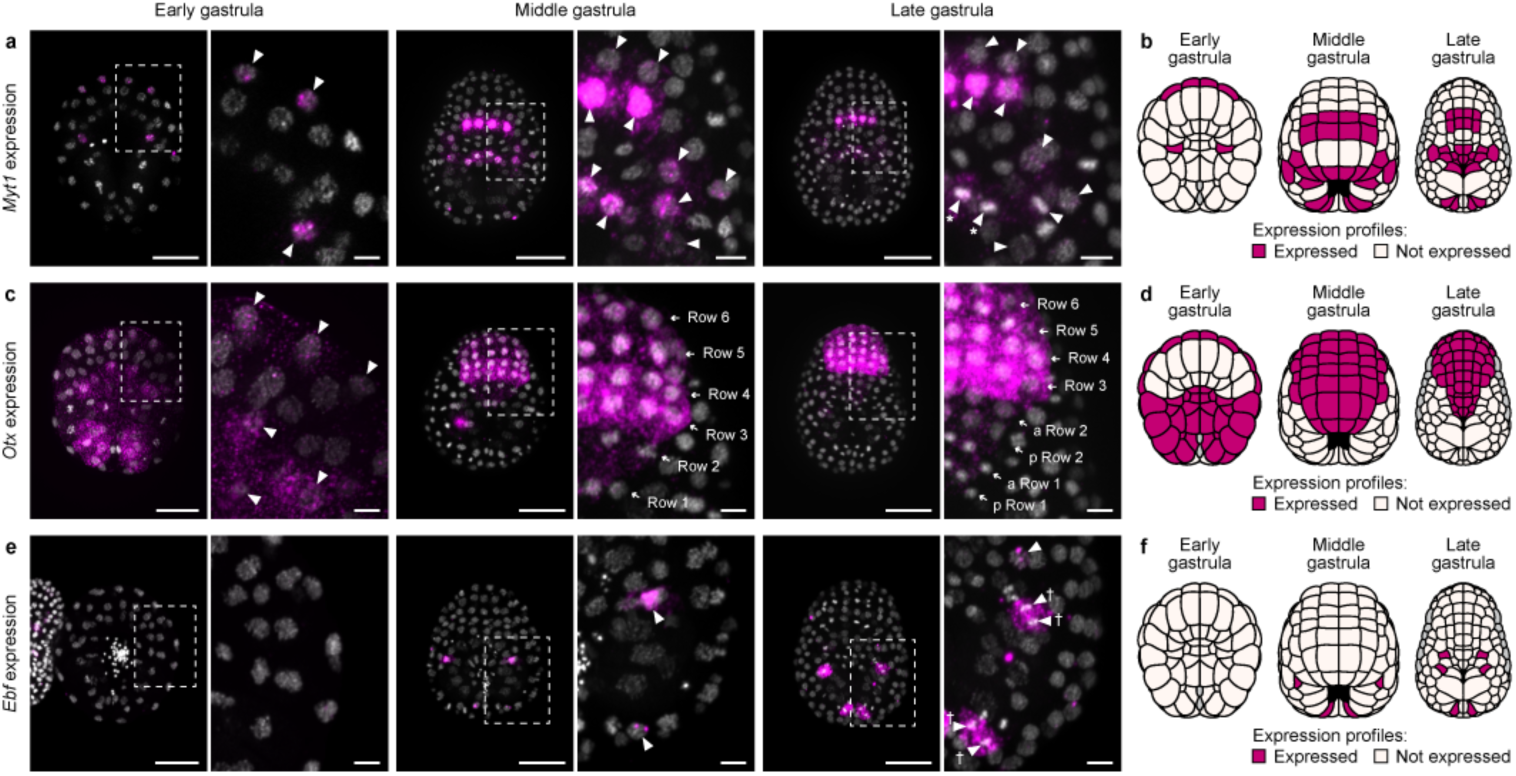
*In situ* hybridization reveals expression patterns of transcription factor genes at single-cell resolution. (**a-h**) Spatial expression patterns of (**a,b**) *Myt1*, (**c,d**) *Otx*, and (**e,f**) *Ebf* at the early, middle, and late gastrula stages. Arrowheads indicate nuclei that express the designated genes. Arrowheads with an asterisk in (**a**) indicate dividing cells that are depicted as two daughter cells in the schematic diagram in (**b**). Arrows in (c) indicate six rows of the neural plate and ANB. At the late gastrula stage, ‘a Row 1’ and ‘p Row 1’ refer to the anterior and posterior daughter cells of row 1 neural plate cells at the middle gastrula stage, respectively; similarly, ‘a Row 2’ and ‘p Row 2’ represent anterior and posterior daughter cells of row 2 cells. Note that epidermal cells anterior to row 6 also express *Otx* in the middle and late gastrula embryos. Arrowheads with daggers denote dividing cells that are illustrated as single mother cells in the schematic in (**f**). Nuclei were stained with DAPI (grey), and *in situ* hybridization signals are colored magenta. Photographs are pseudo-colored z-projections, and the brightness and contrast were linearly adjusted. Scale bar, 50 μm (left panels), 10 µm (right panels). (**b,d,f,h**) Schematic representation of (**b**) *Myt1*, (**d**) *Otx*, and (**f**) *Ebf* expression. Magenta denotes expression. Expression in grey-colored cells was not examined in the present study.

**Figure 3.**
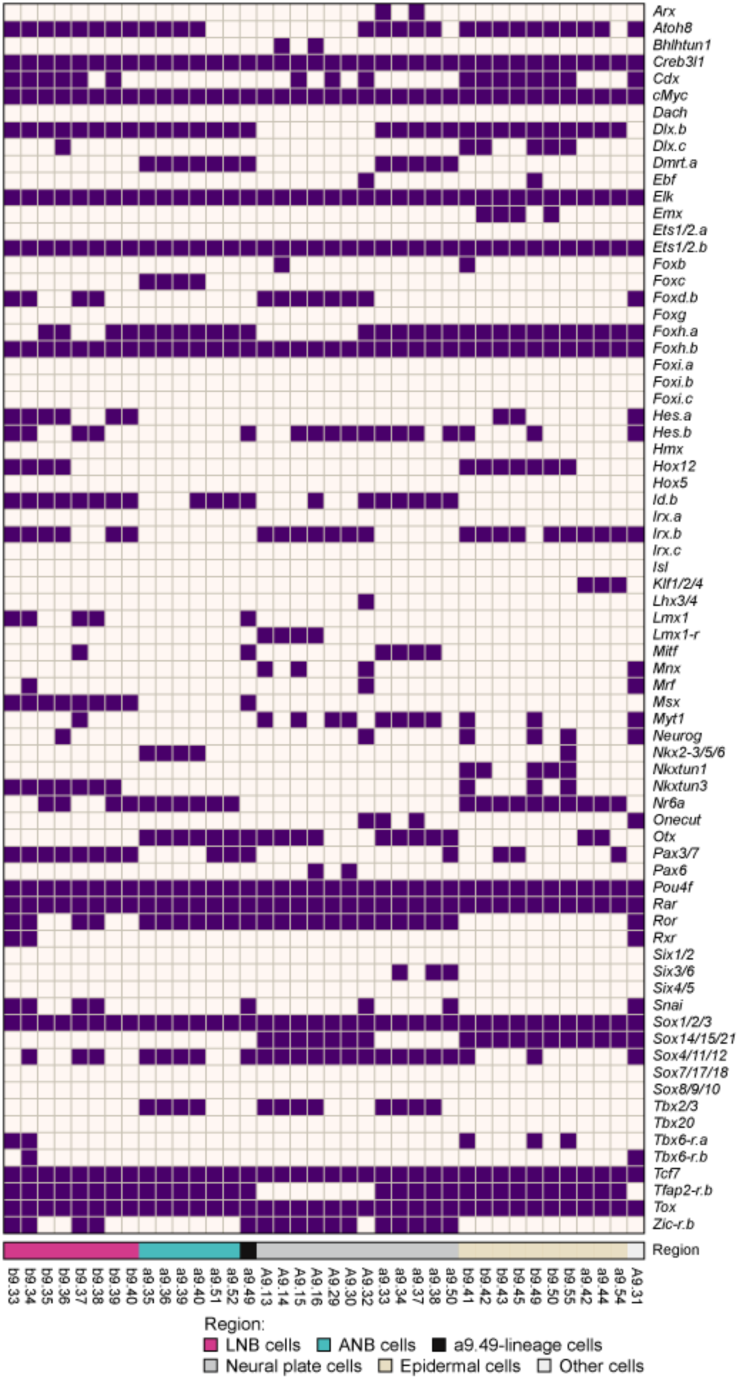
Comprehensive expression profiling of transcription factor genes. The expression matrix of 73 *Ciona* TF genes at the middle gastrula stage. Expression matrices for the other two stages are shown in Supplementary Figure 2. Dark violet denotes expression.

### Three domains in the *Ciona* neural plate border

To characterize cells in the *Ciona* neural plate border, we calculated pairwise expression similarity indices between all possible pairs of cells for each of the three developmental stages (**Fig. 4a-c; Supplementary Table 3**). While all LNB cells of early gastrula embryos (b8.17–b8.20) are grouped into a single cluster with high mutual similarity (**Fig. 4a**), their descendants in middle and late gastrula embryos formed two discrete clusters, b9.33/34/37/38 and b9.35/36/39/40 (or b10.69– 72/77–80) (**Fig. 4b,c**). Cells in the first group laterally abut the neural plate and express *Hes.b*, *Lmx1*, *Snai*, and *Zic-r.b*, while cells in the second group, which are located between the first group and epidermal cells, express *Foxh.a* and *Nr6a* (**see** **Fig. 1**; **Fig. 3, Supplementary Fig. 1,2**). Cells in the first group give rise to ependymal cells of the nerve cord (Nicol & Meinertzhagen, 1988; Nishida, 1987), and cells in the second group give rise to dorsal midline epidermal cells and peripheral sensory neurons (Nishida, 1987; Pasini et al., 2006; Stolfi et al., 2015; Waki et al., 2015). We designated the former LNB cells as “inner lateral neural plate border (iLNB) cells”, and “latter outer lateral neural plate border (oLNB) cells” (**Fig. 4d**).

**Figure 4.**
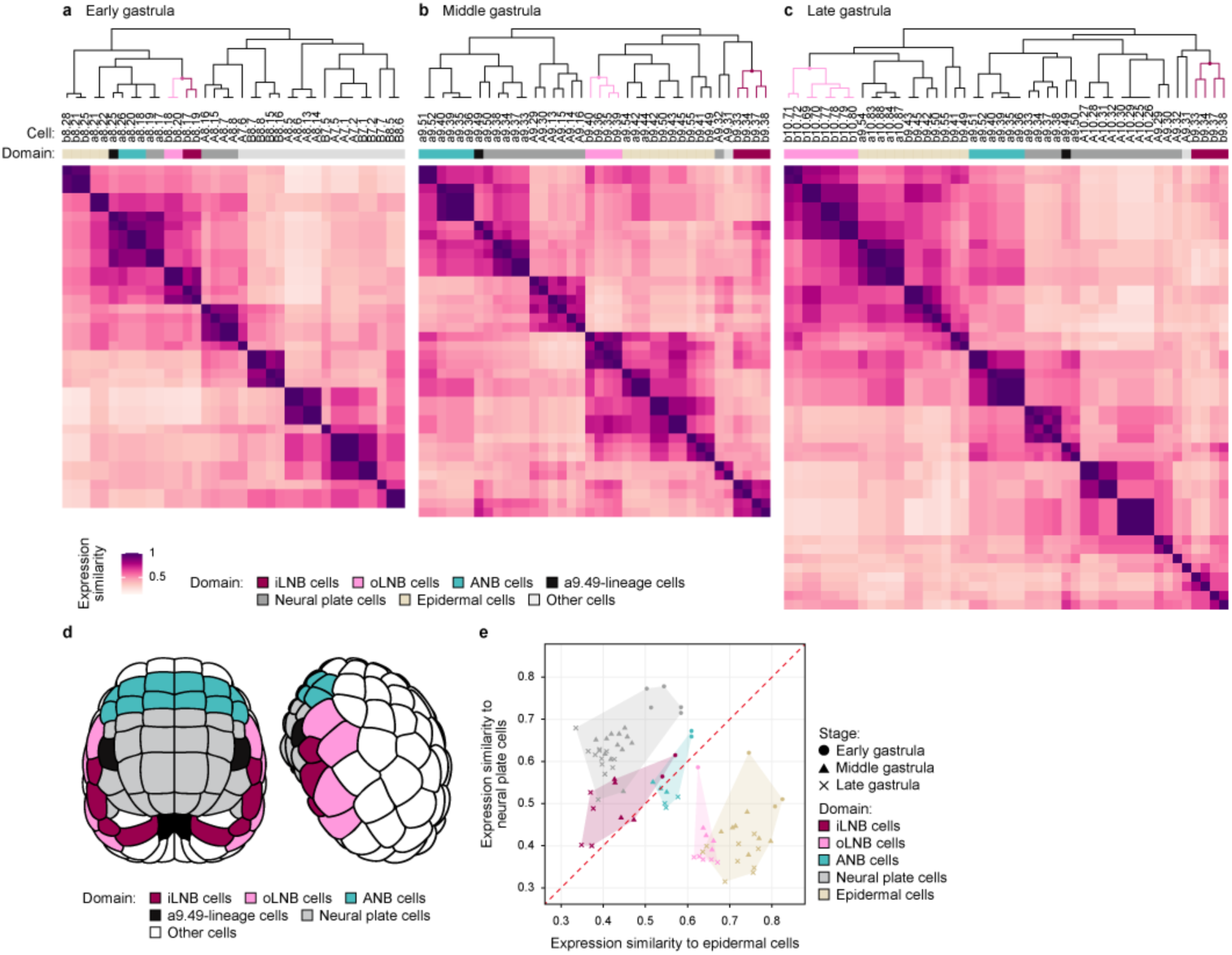
The *Ciona* neural plate border comprises anterior, inner lateral, and outer lateral domains. (**a-c**) Heatmap displaying pairwise gene expression similarity indices during the early (**a**), middle (**b**), and late (**c**) gastrula stages. Similarities were calculated using Jaccard similarity indices, and hierarchical clustering dendrograms were constructed via complete linkage. (**d**) Schematics of the neural plate border and neural plate cells in ascidian embryos at the middle gastrula stages. Cells are colored as follows: iLNB (magenta), oLNB (pink), ANB (cyan), neural plate (grey), and pigment-lineage cells (black). (**e**) Gene expression similarities between individual cells and the entire neural plate or epidermis cell population. Marker shapes indicate developmental stages, while colors indicate the developmental domain. Semi-transparent shaded areas show the convex hull for each domain.

Although ANB cells formed a single cluster in all developmental stages (**Fig. 4a-c**), it was possible to further divide ANB cells into two sub-groups: a sub-group of a8.18 and a8.20 cells and their descendants (a9.35/36/39/40), and a sub-group of a8.26 cells and their descendants (a9.51/52) at all stages (**Fig. 4a-c**). The former cells are located medially and give rise to the oral siphon primordium and papilla cells (Johnson et al., 2024; Liu & Satou, 2019), whereas the latter cells are located laterally and give rise to the anterior trunk epidermal neurons (aATENs) and rostral trunk epidermal neurons (RTENs) (see Figure 1) (Abitua et al., 2015; Deschet & Smith, 2004; Nishida, 1987; Ohtsuka et al., 2014).

Then, we calculated mean expression similarity indices for individual cells against the entire population of either neural plate cells or epidermal cells (**Fig. 4e**). oLNB cells exhibited greater similarity to epidermal cells than to neural plate cells, whereas iLNB cells showed more similarity to neural plate cells than to epidermal cells. For confirmation, we examined expression of a pan-neural marker (*Celf3.a*) (Yagi & Makabe, 2001) and an epidermal marker (*Epib*) (Ueki et al., 1994) by *in situ* hybridization. iLNB cells expressed *Celf3.a*, but not *Epib*, while oLNB cells expressed *Epib*, but not *Celf3.a* (**Supplementary Fig. 3a,b**). In contrast, expression profiles of ANB cells were almost equally similar to those of epidermal and neural plate cells (**Fig. 4e**), and they expressed both *Epib* and *Celf3.a* (**Supplementary Fig. 3a,b**).

### ANB, iLNB, and oLNB cells show transcriptomic similarity to neurogenic placode cells, neural crest/tail bud cells, and median fin fold ectoderm cells of zebrafish embryos, respectively

To identify structures homologous to ascidian ANB, iLNB, and oLNB domains in vertebrate embryos, we analyzed enrichment for zebrafish orthologs of *Ciona* domain-specific gene sets among zebrafish ectodermal cells using publicly available spatial transcriptome datasets from zebrafish embryos 12 h post-fertilization (hpf) (Wan et al., 2026). We first extracted genes expressed differentially among the neural plate, iLNB, oLNB, ANB, and epidermal domains of *Ciona* embryos (hereafter, we denominate these sets as domain-specific gene sets) (**Supplementary Table 4**). We then calculated UCell scores to assess expression enrichment of orthologs of these gene sets in zebrafish spatial transcriptome datasets (Andreatta & Carmona, 2021). Dorsal and transverse views of spatial mapping showed that UCell scores for zebrafish orthologs of neural plate and epidermis domain-specific gene sets were high in cells of the zebrafish central nervous system and epidermis, respectively (**Fig. 5a; Supplementary Fig. 4**). Similarly, the UCell score for the ANB domain-specific gene set was high in the zebrafish neurogenic placode (**Fig. 5a; Supplementary Fig. 4**), and the UCell score for the iLNB domain-specific gene set was high in the zebrafish neural crest and tail bud (**Fig. 5a; Supplementary Fig. 4**). This observation is consistent with our previous finding that iLNB cells have properties of vertebrate neural crest cells and NMPs in the tail bud (Ishida & Satou, 2024).

**Figure 5.**
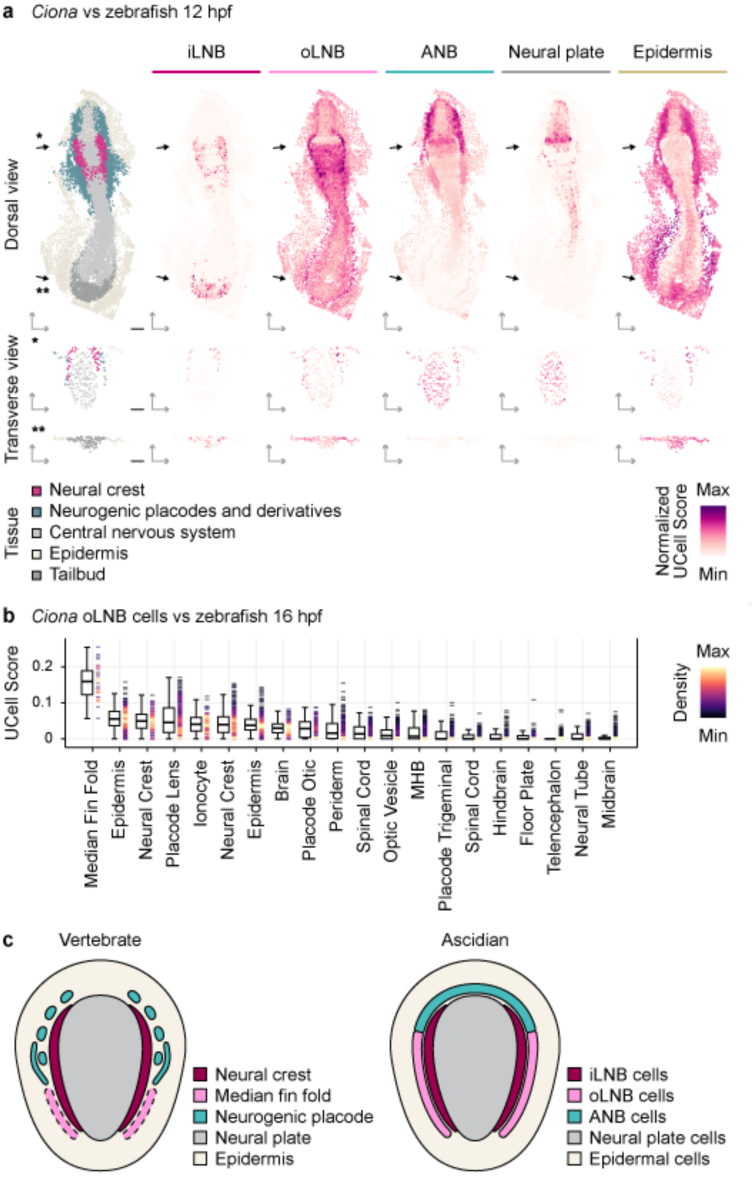
Cross-species comparisons of the *Ciona* iLNB, oLNB, and ANB domains with zebrafish transcriptome data. (**a**) Spatial mapping of UCell scores of zebrafish orthologs for each of the *Ciona* domain-specific gene sets onto the zebrafish spatial transcriptome dataset. Top left: dorsal view of zebrafish embryos with annotated clusters. Top right: UCell scores for zebrafish orthologs of the *Ciona* neural plate, iLNB, oLNB, ANB, and epidermis domain-specific gene sets. Black arrows indicate the anterior-posterior positions of the two transverse views shown at the bottom. (**b**) UCell scores of the *Ciona* oLNB domain-specific gene set in individual clusters of a single-cell transcriptome dataset for zebrafish 16-hpf embryos. Rug plot colors indicate data density. For the box plot, the central line of each box indicates the median, while the lower and upper hinges represent the first (25^th^ percentile) and third (75^th^ percentile) quartiles, respectively. The whiskers extend to the furthest data points that fall within 1.5x the interquartile range from the hinges. (**c**) Schematic model proposing a conserved tripartite spatial organization of the neural plate border across vertebrates and ascidians.

Because the UCell score for the oLNB domain-specific gene set was high in most ectodermal tissues of 12-hpf embryos, we analyzed expression enrichment of the oLNB domain-specific gene set using publicly available single-cell transcriptome datasets of zebrafish 16-hpf embryos (Lange et al., 2024). We found that the UCell score for the oLNB domain-specific gene set was conspicuously high in cluster 21, which strongly expressed *dlx5a*, *prdm1a*, *msx2b*, *sp7*, *sp9*, and *dlx3b* (**Supplementary Fig. 5a-c**). These genes are markers for presumptive median fin fold (MFF) ectoderm (Abe et al., 2007; Akimenko et al., 1994, 1995; Heude et al., 2014; C. Thisse & Thisse, 2005), which is a rudiment of the median fin epidermis and is derived from the posterior lateral neural plate border in zebrafish embryos (Heude et al., 2014; Narboux-Neme et al., 2019). Expression enrichment of orthologs for the oLNB domain-specific gene set is summarized in Figure 4b. This similarity is supported by the observation that oLNB cells that express *Dlx.c*, subsequently underwent a dynamic convergent movement toward the dorsal midline (**Supplementary Fig. 5d**), like zebrafish MFF (Heude et al., 2014).

In zebrafish embryos, both neural crest cells and Rohon-Beard neurons are differentiated in the posterior neural plate border (Artinger et al., 1999; Cornell & Eisen, 2002). To investigate spatial organization of the zebrafish neural plate border, we analyzed spatial expression patterns of markers for neural crest, Rohon-Beard neurons, and MFF ectoderm. We found that the UCell score for genes that were identified as differentially expressed in presumptive MFF ectoderm in 16-hpf embryos was high in the posterior lateral neural plate border in zebrafish 12-hpf embryos (**Supplementary Fig. 6a**). Notably, MFF markers were strongly expressed in cells lateral to cells strongly expressing neural crest markers and Rohon-Beard neuron markers (**Supplementary Fig. 6a-d**). This clear spatial segregation indicates that MFF ectoderm originates from a lateral domain in the neural plate border, distinct from regions that give rise to the neural crest and Rohon-Beard neurons. Thus, the zebrafish neural plate has a mediolateral spatial organization similar to that of the ascidian neural plate (**Fig. 5c**).

## Discussion

Because sets of regulatory factors define cell types (Arendt et al., 2016; Davidson et al., 2003), the unbiased gene expression dataset in the present study allowed us to define three transcriptomically distinct domains, or domains with three distinct “regulatory states,” in the *Ciona* neural plate border region (ANB, iLNB, and oLNB domains). These regulatory states were similar to those of zebrafish neurogenic placode cells, neural crest/tail bud cells, and the presumptive MFF ectoderm.

## Anterior neural plate border cells

Our data suggest that ANB cells can be subdivided into two distinct subpopulations: cells in the medial subdomain that give rise to the oral siphon primordium and papilla cells (Johnson et al., 2024; Liu & Satou, 2019; Roure et al., 2025), and cells in the lateral subdomain that give rise to epidermal cells and sensory neurons (Abitua et al., 2015; Deschet & Smith, 2004; Nishida, 1987; Ohtsuka et al., 2014). In addition, the signalling pathway governing specification is distinct between the former and latter cells (Hudson & Yasuo, 2005; Liu et al., 2023; Liu & Satou, 2019; Ohtsuka et al., 2014; Wagner & Levine, 2012). Thus, the last common ancestor of vertebrates and tunicates may have had multiple groups of neurogenic placode-like cells.

Previous studies suggested that *Ciona* ANB cells resemble vertebrate olfactory placode cells (Abitua et al., 2015). Indeed, *Emx*, *Foxg*, *Six1/2*, *Six3/6*, and *Sp.b* (*Zf220*), which are expressed in vertebrate neurogenic placodes, initiate their expression in ANB cells later (Abitua et al., 2015; Johnson et al., 2024; Liu & Satou, 2019), although they were not expressed in early embryos that we examined. Comprehensive gene expression assays using later embryos may reveal vertebrate homologs of ascidian ANB cells.

## Inner lateral neural plate border cells

*Ciona* iLNB cells have properties resembling those of vertebrate neural crest cells and NMPs (Ishida & Satou, 2024). Consistent with this previous study, our present study showed that *Ciona* iLNB cells are similar in their regulatory state to both neural crest and tail bud cells of zebrafish. Indeed, iLNB cells express orthologs of TF genes specifying the neural plate border and neural crest cells in vertebrates, which include *Dlx.b*, *Msx*, *Snai*, *Zic-r.b*, *Tfap2-r.b*, *Pax3/7*, *Ets1/2.b*, *Lmx1*, *Foxd.b*, *Mitf* and *Id.b* (Martik & Bronner, 2017), and those expressed in tail bud cells in vertebrates, which include *Msx*, *Foxd.b*, *Hes.b*, *Rxr*, *Cdx*, *Hox12*, and *Tbx6-r.a/Tbx6-r.b* (Connors et al., 1999; Schmid et al., 2000; B. Thisse & Thisse, 2004; C. Thisse & Thisse, 2005, 2008).

Interestingly, pigment-lineage cells (a9.49-lineage cells), which are also likely homologous to vertebrate neural crest cells (Abitua et al., 2012; Fatieieva et al., 2025; Todorov et al., 2024), did not express the tail bud markers, *Rxr*, *Cdx*, *Hox12*, or *Tbx6-r.a/Tbx6-r.b*, while they did express the neural crest markers, *Dlx.b*, *Msx*, *Snai*, *Zic-r.b*, *Tfap2-r.b*, *Pax3/7*, *Ets1/2.b*, *Lmx1*, *Mitf*, and *Id.b*. Pigment-lineage cells and iLNB cells are derived from different cell lineages. Pigment-lineage cells develop from the anterior animal cells (a4.2) of 8-cell embryos, and iLNB cells develop from the posterior animal cells (b4.2) (Nishida, 1987; Nishida & Satoh, 1983). It is possible that these ascidian pigment-lineage neural-crest-like cells evolved independently from iLNB cells.

## Outer lateral neural plate border cells

oLNB cells are thought to be evolutionarily homologous to vertebrate neural crest cells (Horie et al., 2018; Stolfi et al., 2015; Waki et al., 2015), neurogenic placode cells (Papadogiannis et al., 2022), or dorsal median fin ectoderm (Pasini et al., 2006). Our observations in the present study do not strongly support similarities of ascidian oLNB cells to either vertebrate neurogenic placode or neural crest cells. Indeed, oLNB cells did not express neurogenic placode markers, *Six1/2*, *Six3/6*, *Six4/5*, *Eya*, *Dach*, or *Foxg*, nor an anterior marker, *Otx*, even at later stages (Hudson & Lemaire, 2001; Imai et al., 2004; Liu & Satou, 2019; Mazet et al., 2005). Instead, they expressed posterior markers, *Cdx* and *Hox12*, orthologs of which are not expressed in neurogenic placodes in vertebrate embryos (B. Thisse et al., 2004). Furthermore, these cells showed gene expression profiles distinct from those of iLNB cells, and did not express neural crest markers, *Snai*, or *Lmx1*.

Instead, our cross-species comparison revealed significant similarity of *Ciona* oLNB cells with the zebrafish posterior lateral neural plate border and its derivative, MFF (**Fig. 4b; Supplementary Fig. 5**). During zebrafish development, MFF ectoderm arises from a posterior neural plate border domain expressing *dlx5a* (Heude et al., 2014; Narboux-Neme et al., 2019). During neurulation, *dlx5a*-expressing neural plate border cells follow a dynamic convergent movement toward the dorsal midline to form the presumptive MFF ectoderm (Heude et al., 2014; Narboux-Neme et al., 2019). In *Ciona* embryos, descendants of oLNB cells similarly undergo a dynamic convergent movement toward the dorsal midline and form the dorsal fin (Nishida, 1987; Pasini et al., 2006). Therefore, it is highly likely that *Ciona* oLNB cells are homologous to vertebrate neural plate border cells that give rise to MFF ectoderm, which is suggested to be an ancestral trait of chordates (Mabee et al., 2002; Mussini et al., 2024; Pasini et al., 2006).

The *Ciona* dorsal midline is a region where not just the dorsal median fin forms, but also a region from which caudal epidermal sensory neurons (CESNs) and bipolar tail neurons (BTNs) differentiate (Lanoizelet et al., 2024; Pasini et al., 2006; Stolfi et al., 2015; Waki et al., 2015). However, in vertebrate embryos, neurons are not known to arise from MFF ectoderm. Two possible scenarios may explain this. In the first scenario, vertebrates lost ancestral neurogenic potential in the MFF ectoderm (Pasini et al., 2006). In the second scenario, neurogenic potential was acquired in the ascidian lineage after divergence from the vertebrate lineage. We have previously presented evidence indicating that the gene circuit for specifying the dorsal CESNs of ascidian embryos emerged via a co-option of the *Msx*-driven regulatory circuit that specifies the ventral CESNs (Waki et al., 2015). If the first scenario is correct, neurogenic potential may have been lost in ascidian iLNB and vertebrate MFF ectoderm. If the second explanation is correct, this co-option likely occurred in the tunicate lineage after its divergence from the vertebrate lineage, and vertebrates somehow recruited or innovated a gene circuit that produce peripheral neurons derived from the neural crest.

## The neural plate border of embryos of the last common ancestor of vertebrates and ascidians likely had the tripartite organization

In vertebrate embryos, neural crest and neurogenic placodes give rise to cranial structures, including special sensory organs, cranial ganglia, and jaws. Our findings reveal that ascidian embryos possess a tripartite neural plate border structurally similar to that of zebrafish embryos. Therefore, this organization was likely present in the last common ancestor of vertebrates and tunicates. The tripartite neural plate border may have functioned as an ancestral blueprint, elaboration and sophistication of which in the vertebrate lineage after its divergence from ascidians led to evolutionary emergence of the “new head” (Gans & Northcutt, 1983) and median fins.

## Acknowledgements

We thank Gembu Abe, Yi-Hsien Su and Jr-Kai Yu for critical comments on the manuscript; Yasushi Okochi for his advice about the single-cell transcriptome analysis. We also thank Reiji Masuda, Chikako Imaizumi (Kyoto University), Toru Miura, Manabu Yoshida (University of Tokyo), and other members working under the National BioResource Project for *Ciona* (MEXT, Japan) at Kyoto University and the University of Tokyo for providing experimental animals.

This research was supported by a GINPU grant from Kyoto University to TI, grants from the Japan Society for the Promotion of Science under the grant numbers 25KJ1652 to TI, 21H02486/21H05239/24K02032/24K21274 to YS.

## Author contributions

TI and YS conceived and designed experiments. TI performed experiments. TI and YS analyzed data. TI wrote the original draft of the manuscript; YS reviewed and edited the manuscript.

## Data availability

Original image data underlying Figure 2 and Supplementary Figure 1 are available in The Kyoto University Research Information Repository (KURENAI, http://hdl.handle.net/2433/302290; DOI: 10.57723/302290).

## Materials and Methods

### Animals

Adult specimens of *Ciona robusta* (formerly called *C. intestinalis* type A) were obtained from the National BioResource Project for *Ciona* (Satou et al., 2026). For experiments, eggs and sperm were collected from oviducts and sperm ducts of adults. Unfertilized eggs were treated with dechorionation solution [1% Sodium Mercaptoacetate (FUJIFILM, #194-03551), 0.5% Actinase E (KAKEN, #650133), 36 μM NaOH in seawater] to remove the chorion. Dechorionated eggs were washed three times with fresh seawater. Both fertilized eggs and embryos were raised at 18°C.

### Gene identifiers

Identifiers (Satou et al., 2022) for the epidermal marker *Epib* and the pan-neural marker *Celf3.a* are KY21.Chr7.872 and KY21.Chr6.58, respectively. Identifiers for the transcription factor genes examined in this study are listed in Supplementary Table 1.

### Whole-mount fluorescence *in situ* hybridization

cDNAs for *Emx*, *Foxg, Hmx*, and *Pax3/7* were amplified from a cDNA library. Primers used for cDNA amplification were as follows: for *Emx*, 5’-CGTCATAGACGCTTGCGTTA-3’ and 5’-CTGAATAGCCGTTTCGCGTT-3’; for *Foxg*, 5’-CAATTGCGATGATGACGCAA-3’ and 5’-ATGACGAACGACGCGGC-3’ ; for *Hmx*, 5’-TCAAACCACGGGATTCCC-3’ and 5’-CTATGACGTCACTGTGCCAA-3’; for *Pax3/7*, 5’-TACAAGCTGCACTACAAGAC-3’ and 5’-GTTGTAAAGTAGTCTGACTTCTG-3’. These fragments were subsequently cloned into the pBluescript II SK(-) vector using an In-Fusion HD Cloning Kit (Takara, #639649). cDNA clones for the remaining genes were obtained from the EST clone collection (Satou et al., 2005). Digoxygenin (DIG) and fluorescein-labeled antisense RNA probes were synthesized using T7 RNA Polymerase (Invitrogen, #18033-019), DIG RNA Labeling Mix (Merck, #11277073910), and Fluorescein RNA Labeling Mix (Merck, #11685619910), and then used for *in situ* hybridization.

Embryos were fixed in 4% paraformaldehyde (PFA) in 0.1 M MOPS buffer (pH 7.5) and 0.5 M NaCl at 4 °C for over 16 h. After washing with 80% ethanol and phosphate-buffered saline containing 0.1% Tween 20 (PBST), embryos were incubated in 3% H_2_O_2_ for 60 min and then washed again with PBST. Then, embryos were treated with 2 μg/mL of proteinase K in PBST for 30 min at 37 °C and washed with PBST. Embryos were again fixed with 4% paraformaldehyde in PBST for 1 h at room temperature, washed with PBST, and immersed in hybridization buffer for at least 1 h at 50 °C. Then, the hybridization buffer was replaced with fresh hybridization buffer containing a DIG- and/or a fluorescein-labeled probe, and embryos were incubated at 50 °C for at least 16 h. Hybridization buffer contained 50% formamide, 5× SSC, 100 μg/mL of yeast transfer RNA, 5× Denhart’s solution, and 0.1% Tween 20. After hybridization, embryos were washed twice at 50 °C for 15 min in 2× SSC, 50% formamide, and 0.1% Tween 20. Embryos were treated for 30 min at 37 °C with 20 μg/mL of RNase A in 10 mM Tris buffer (pH 8.0), 0.5 M NaCl, 5 mM EDTA, and 0.1% Tween 20. Embryos were further washed twice at 50 °C for 15 min in 0.5× SSC, 50% formamide, and 0.1% Tween 20, and the washing solution was replaced with PBST. Embryos were blocked at room temperature for 60 min with 0.1% BSA in PBST and then exposed to mouse anti-digoxigenin (Merck, #11333062910; 1:100 dilution) or rabbit anti-fluorescein (Abcam, #ab19491; 1:1,000 dilution) diluted in Can Get Signal Immunostain Immunoreaction Enhancer Solution A (Toyobo, #NKB-501) at 4 °C overnight. After washing with PBST, embryos were incubated with an HRP-conjugated secondary antibody (Thermo Fisher, B40961 and B40922; 1:1 dilution). Embryos were washed with PBST and TNT (100 mM Tris buffer (pH 7.5), 150 mM NaCl, and 0.1% Tween 20). Then, embryos were stained with Alexa Fluor 488 or 555 tyramide reagent (Thermo Fisher, #B40953 and #B40955). Before observation with a microscope, embryos were treated with a TrueVIEW Autofluorescence Quenching Kit (Vector Laboratories, #SP-8400-15).

Gene expression profiles were described for neural plate, neural plate border, and epidermal cells. For the epidermal lineage, the analysis was confined exclusively to dorsal cells proximal to the neural plate and neural plate border (early gastrula: a8.21, a8.22, b8.21, b8.25, b8.28; middle gastrula: a9.42, a9.44, a9.54, b9.41, b9.42, b9.43, b9.45, b9.49, b9.50, b9.55; late gastrula: a10.83, a10.84, a10.87, a10.88, a9.54, b9.41, b9.42, b9.43, b9.45, b9.49, b9.50, b9.55). We used previously published expression data for the following genes: *Dlx.b*, *Ets1/2.b*, *Id.b*, *Lmx1*, *Msx*, *Pax3/7*, *Tfap2-r.b*, and *Zic-r.b* at the early and middle gastrula stages, and *Snai* at the early, middle, and late gastrula stages (Ishida & Satou, 2024).

## Data analysis

### Identification of candidate Ciona genes

To identify TF genes expressed in the neural plate, neural plate border, and adjacent non-neural ectodermal cells at the early to late gastrula stages, published single-cell transcriptome data for *Ciona* initial and middle gastrula embryos were used (Cao et al., 2019) (SRA accession numbers: SRR9050985 to SRR9050988). Data were mapped to the latest genome assembly (HT version) (Satou et al., 2019) and the latest version of the gene model set (KY21 version) (Satou et al., 2022) using Cell Ranger software (v.6.12, 10x Genomics). Cells were annotated using Loupe Browser (10x Genomics) based on expression of specifically enriched genes that are expressed in specific cells. Identified clusters are listed in Supplementary Table 5.

Of the 303 TF genes listed in the Ghost Database (Satou et al., 2005), 58 TF genes were enriched (average expression > 1 and *p*-value < 0.05) in at least one cluster identified above (**Supplementary Fig. 2**; **Supplementary Table 1**) against all other cells. Because abundant maternal RNA prevents detection of zygotic gene transcripts in *in situ* hybridization, 11 TF genes with abundant maternal expression (TPM>50) were excluded using publicly available bulk RNA-sequencing datasets from *Ciona* unfertilized eggs (Frese et al., 2024) (SRA accession numbers: SRR25223655 and SRR25223647). The excluded 11 genes are *Hmgb1/2*, *Mxd*, *Ets1/2.b*, *Smad1/5/9*, *Tcf3*, *Creb3l1*, *Erf.b*, *Foxtun1*, *Rar*, *Ror*, and *Ybx* (**Supplementary Table 1**). As a result, the initial list contained 47 genes. In addition, *Arx*, *Emx*, *Foxc*, *Foxg*, *Hmx*, *Isl*, *Lhx3/4*, *Mitf*, *Mrf*, *Nkx2-3/5/6*, *Pax3/7*, *Pou4f*, *Six1/2*, *Six3/6*, *Sox7/17/18*, *Tbx6-r.b*, and *Tox*, which are expressed in these cells (Abitua et al., 2012; Coulcher et al., 2020; Ikeda et al., 2013; Imai et al., 2004, 2006; Liu & Satou, 2019; Papadogiannis et al., 2022; Wagner & Levine, 2012; Waki et al., 2015) were also re-examined (**Supplementary Table 1**). Similarly, *Arx*, *Dach*, *Emx*, *Foxc*, *Foxd.b*, *Foxg*, *Foxi.a*, *Foxi.b*, *Foxi.c*, *Hmx*, *Irx.a*, *Irx.c*, *Isl*, *Mitf*, *Myc*, *Pax3/7*, *Pou4f*, *Rxr*, *Six1/2*, *Six3/6*, *Six4/5*, *Sox8/9/10*, and *Tbx20*, which are *Ciona* orthologs for vertebrate TF genes expressed in the neural plate border, neural crest, or neurogenic placodes (Betancur et al., 2010; Gómez-Skarmeta et al., 2003; Simões-Costa & Bronner, 2015; Thawani & Groves, 2020; B. Thisse et al., 2004), were examined (**Supplementary Table 1**).

### Calculation of pairwise expression similarities

Pairwise similarity calculations and hierarchical clustering were performed using an expression matrix derived from *in situ* hybridization data and R software. Gene expression data were first binarized (1 for expression; 0 for no expression). Jaccard similarity indices (Jaccard, 1901, 1912) between all possible cell pairs in individual stages were calculated to compare pairwise cell similarity. Hierarchical clustering dendrograms were constructed using the complete linkage algorithm. To evaluate similarities of gene expression profiles between individual cells and the entire populations of neural plate cells or epidermal cells, average Jaccard similarity indices for each cell against the entire population of neural plate cells or epidermal cells during the same developmental stage were calculated.

### Cross-species transcriptome analysis

For a cross-species comparison between *Ciona in situ* hybridization data and zebrafish spatial (Wan et al., 2026) and single-cell (Lange et al., 2024) transcriptome data, expression profiles of zebrafish orthologs for *Ciona* TF genes differentially expressed in the neural plate, iLNB, oLNB, ANB, and epidermal cells were examined. To identify *Ciona* DEGs for each domain at the middle to late gastrula stages, an expression matrix was prepared from *in situ* hybridization data during the middle and late gastrula stages. Two-sided Fisher’s exact tests were performed to compare the proportion of cells that expressed a gene of interest in a target domain with an average for the remaining domains. *P*-values were adjusted for multiple testing using the Benjamini-Hochberg method (Benjamini & Hochberg, 1995). Genes with adjusted *p*-values (FDR) < 0.05 were considered statistically significant (**Supplementary Table 4**). These DEG sets comprised both positive and negative markers, defined by positive and negative log_2_ odds ratios, respectively.

Processed spatial transcriptome datasets for zebrafish 12-hpf were downloaded from Dryad (https://datadryad.org/dataset/doi:10.5061/dryad.j0zpc86v9; weMERFISH_combined_C_6s_E1_rescaled_z.h5ad and weMERFISH_combined_C_6s_E2_rescaled_z.h5ad). Data were analyzed using R (v.4.5.1) with the Seurat package (v.5.4.1). Spatial transcriptomic data in .h5ad format were imported and converted to Seurat objects. For downstream analyses, cells were filtered to retain only those annotated as “Ectoderm” in ‘germlayer’ metadata using the ‘subset’ function of Seurat. Then, expression scores for *Ciona* DEG sets of each domain were calculated using the UCell package (v.2.12.0) for each individual cell.

To evaluate gene expression along the mediolateral axis at different anteroposterior coordinates, spatial coordinates of cells were transformed into a normalized mediolateral axis. Spatial coordinates of cells were projected onto defined reference lines to determine their anteroposterior and mediolateral positions. Next, lateral boundaries at each anteroposterior position were defined by calculating the minimum and maximum mediolateral coordinates of cells annotated as neural tissues (‘Diencephalon’, ‘Floor Plate’, ‘Forebrain Ventral’, ‘Hindbrain Neurons’, ‘Hindbrain R1+2’, ‘Hindbrain R3+4’, ‘Hindbrain R5+6’, ‘Hindbrain R7’, ‘Midbrain’, ‘Optic Cup’, ‘Spinal Cord’, ‘Spinal Cord Differentiated’, and ‘Tailbud’). These boundaries were then smoothed using LOESS regression. Finally, a midline was calculated as the midpoint between the smoothed left and right boundaries, and the mediolateral position of each cell was normalized such that the midline was defined as 0, and the left and right boundaries were defined as -1 and +1, respectively.

Processed single-cell transcriptome data for zebrafish 16-hpf were downloaded from ZebraHub (https://zebrahub.sf.czbiohub.org/; zf_atlas_16hpf_v4_release.h5ad) (Lange et al., 2024). Data were analyzed using R (v.4.5.1) with the Seurat package (v.5.4.1). Single-cell transcriptomic data in .h5ad format were imported and converted to Seurat objects. Cells were first filtered to retain only those annotated with ectoderm-derived tissues based on ‘zebrafish_anatomy_ontology_class’ metadata (‘brain’, ‘ectodermal cell’, ‘floor plate’, ‘lens placode’, ‘midbrain hindbrain boundary’, ‘neural crest’, ‘neural tube’, ‘optic vesicle’, ‘otic placode’, ‘periderm’, ‘spinal cord neural tube’, ‘telencephalon’, ‘trigeminal placode’). 2,000 highly variable genes were identified using the ‘FindVariableFeatures’ function of Seurat. Data were scaled using the ‘ScaleData’ function of Seurat, and principal component analysis (PCA) was performed using the ‘RunPCA’ function of Seurat. To cluster cells, the ‘FindNeighbors’ function was used with the parameter ‘dims = 1:30’, followed by the ‘FindClusters’ function using the Leiden algorithm with ‘resolution = 1.8’. Dimensionality reduction was performed using the ‘RunUMAP’ function with the parameter ‘dims = 1:30’. Differentially expressed genes were identified using the ‘FindAllMarkers’ function with the parameters ‘logfc.threshold = 2’ and ‘min.pct = 0.25’ and filtered for an FDR < 0.01. The MFF ectoderm-specific gene set was defined as the top 50 differentially expressed genes ranked by FDR and log2 fold-change (Supplementary Table 6). Expression scores for the *Ciona* DEG sets of each domain were calculated using the UCell package (v.2.12.0) for each cell.

---

**Supplementary Fig. 1.**
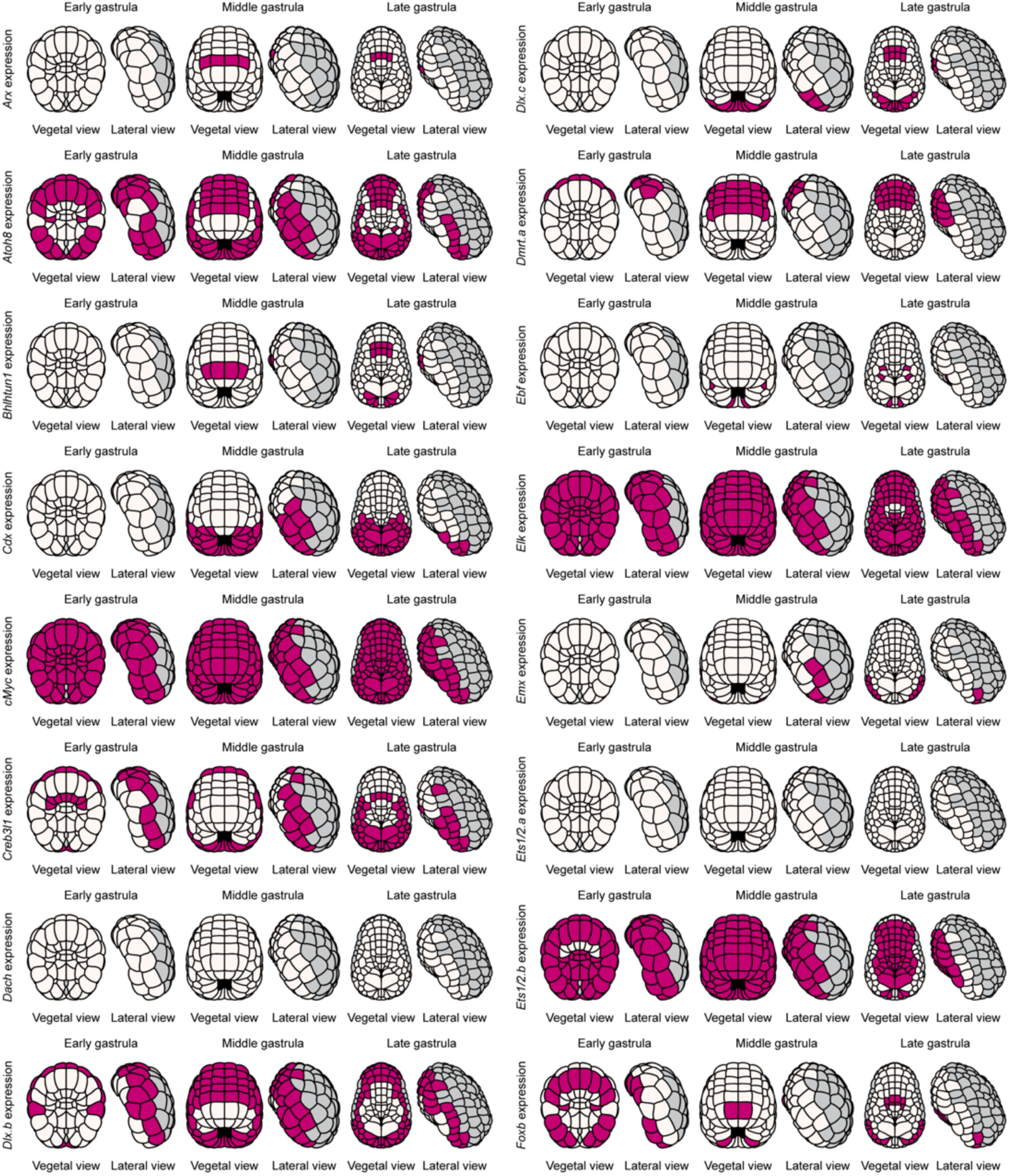
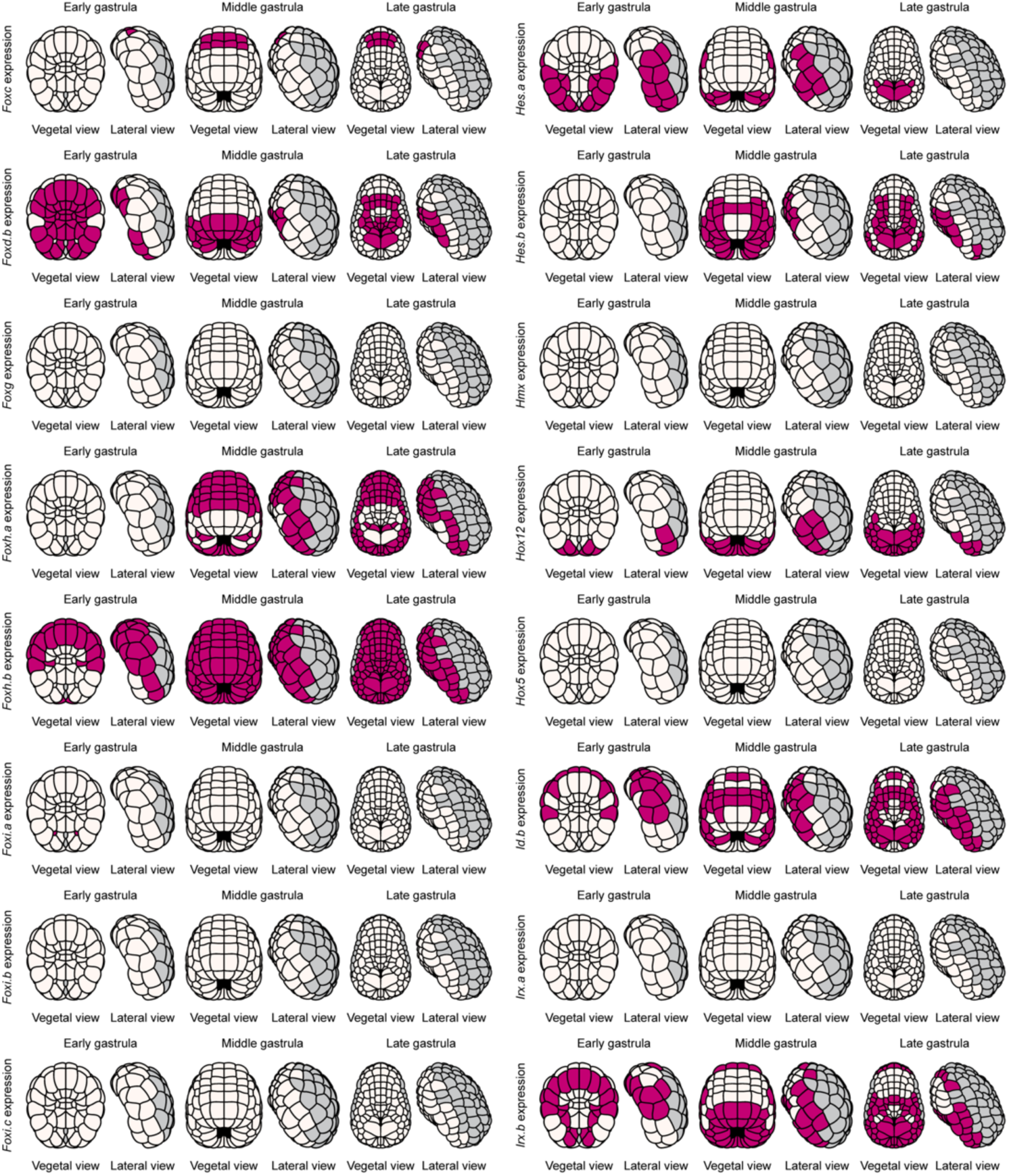
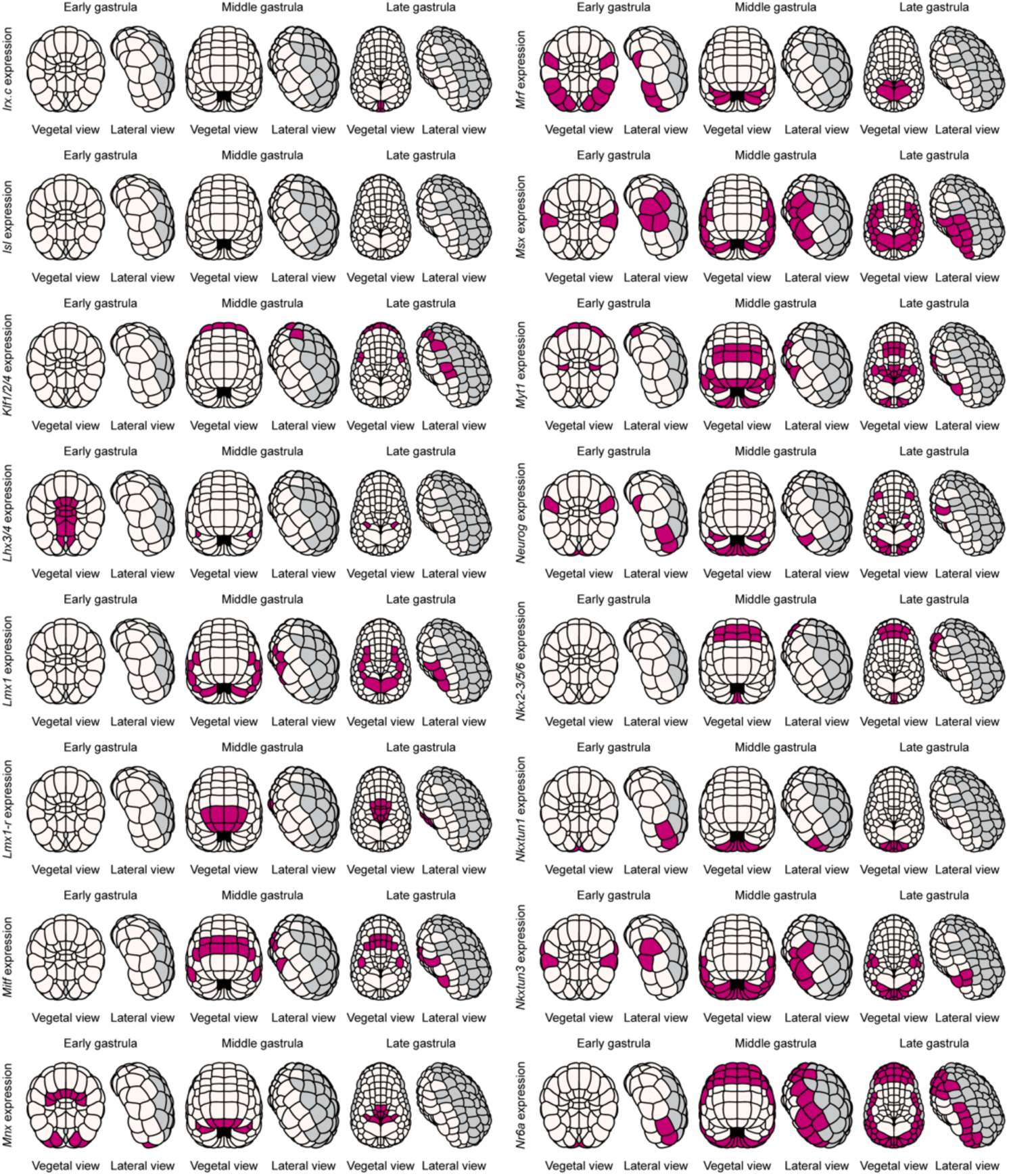
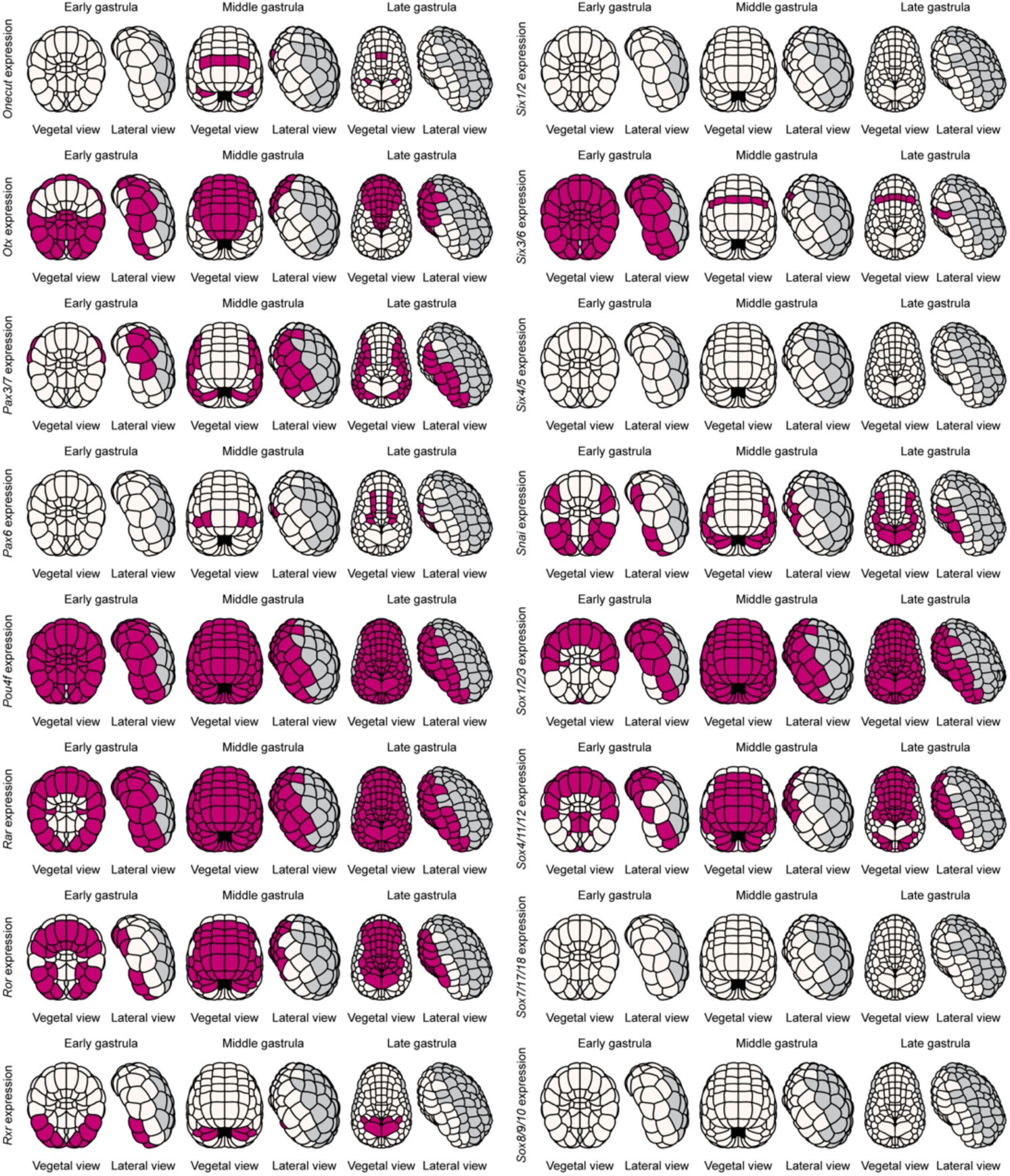
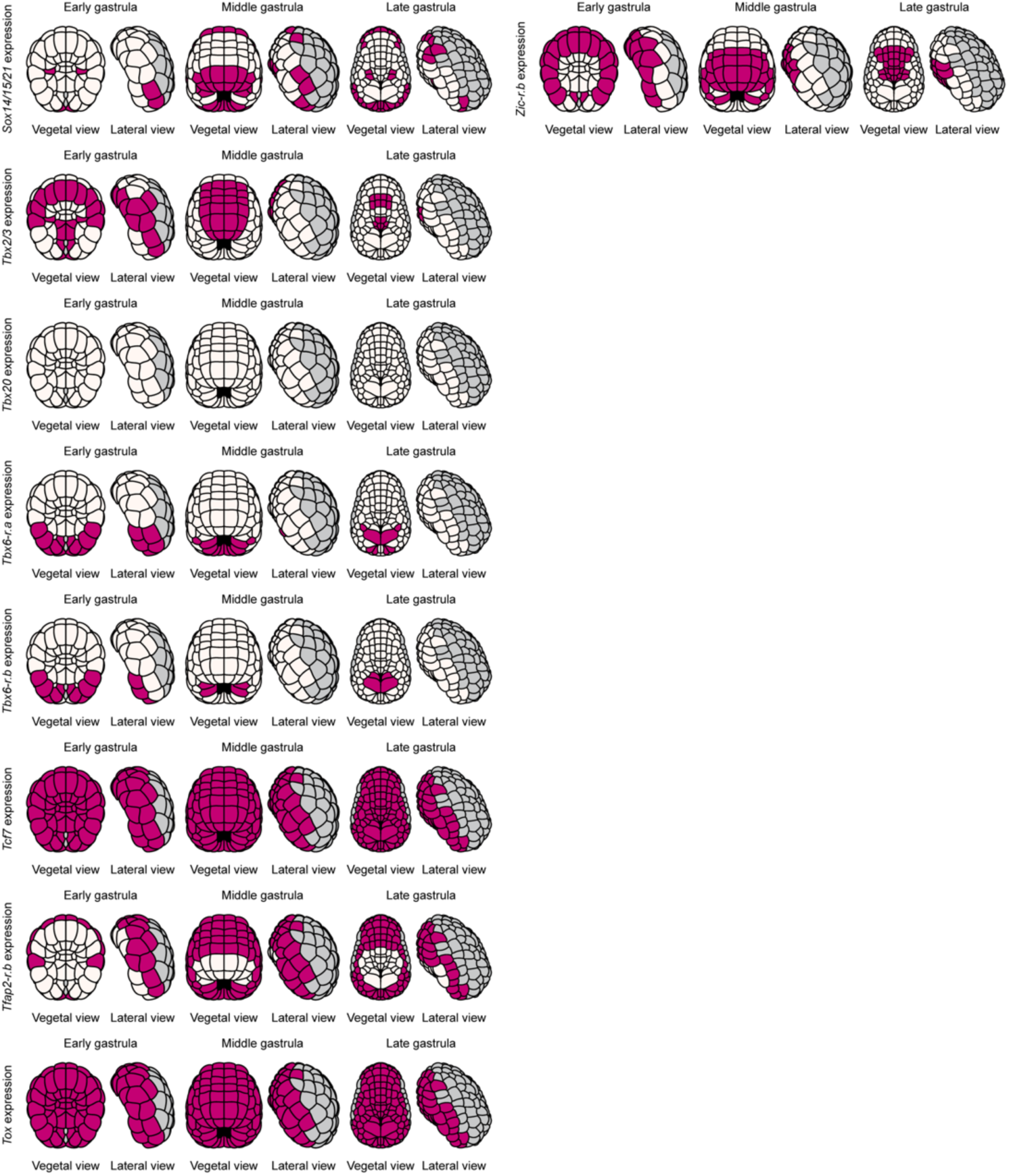
Expression patterns of *Ciona* 73 TF genes. Spatial expression and schematic representation of 73 *Ciona* TF genes during the early (eG), middle (mG), and late (lG) gastrula stages. Nuclei were stained with DAPI (grey), and *in situ* hybridization signals are colored magenta. Photographs are pseudo-colored z-projections. Scale bar, 50 μm. In schematics, dark violet denotes cells with expression, and grey indicates cells that we did not examine. Brightness and contrast of photographs were linearly adjusted.

**Supplementary Fig. 2.**
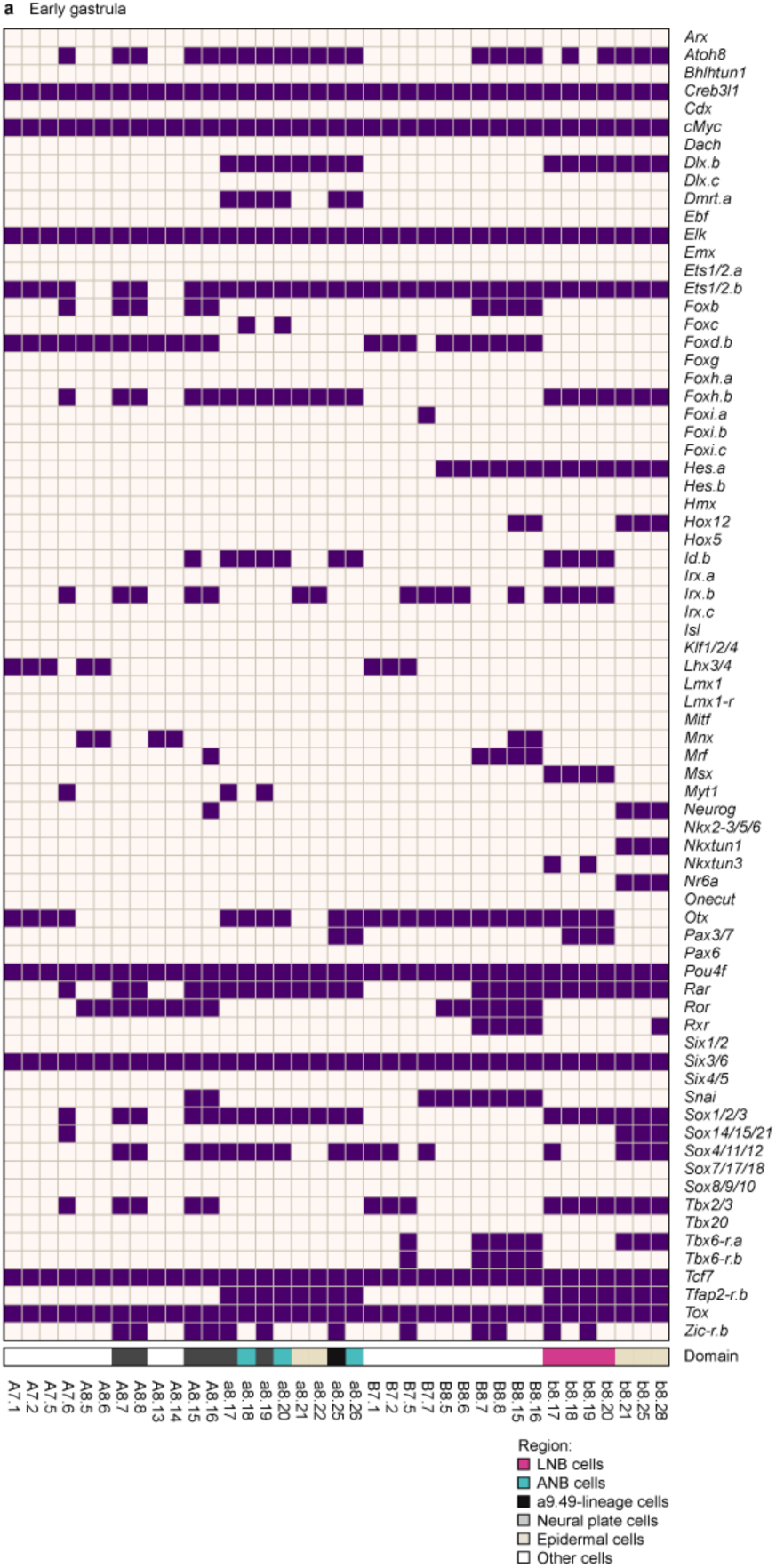
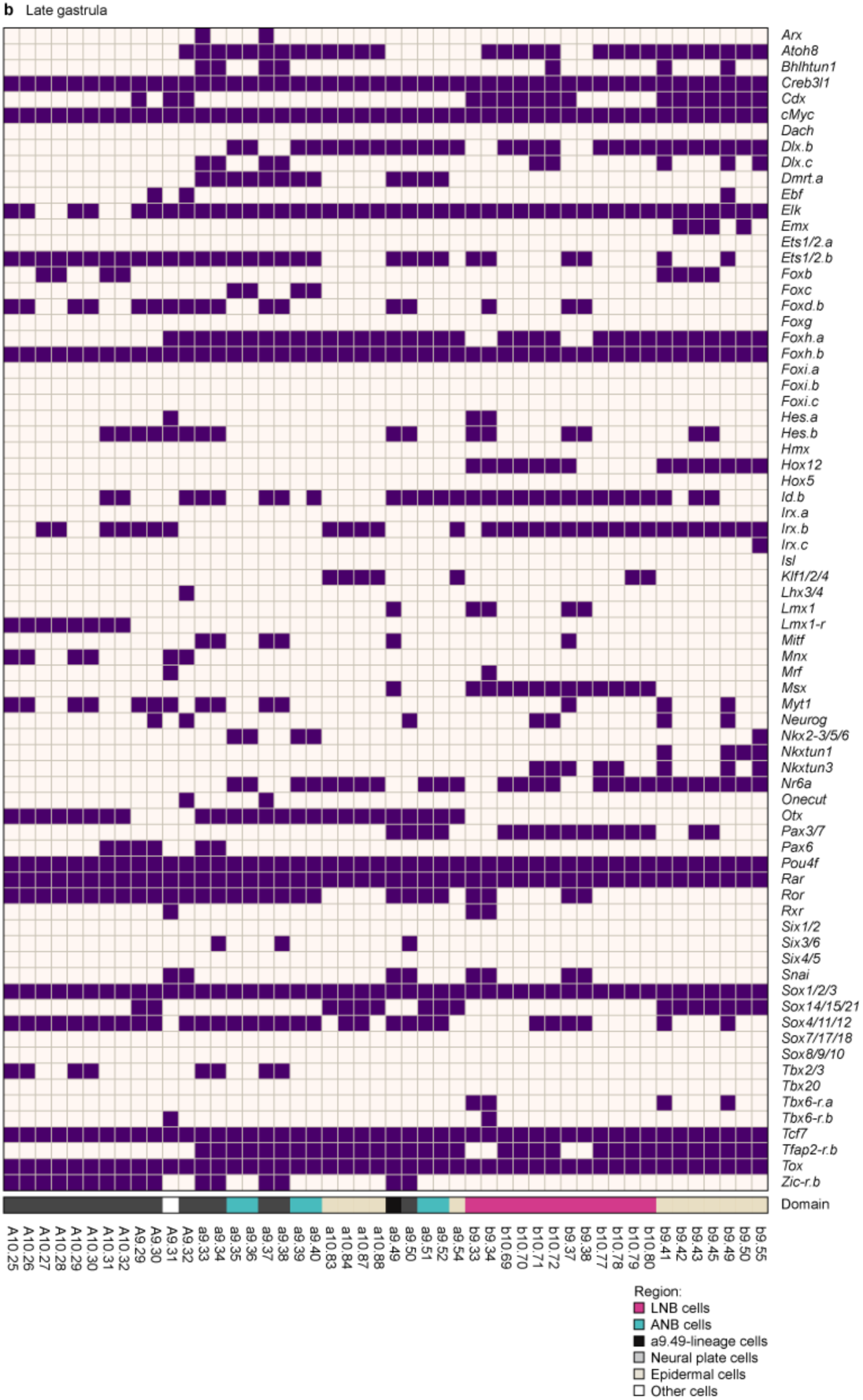
Detailed expression patterns of 73 TF genes shown in. Fig. 2. (**a,b**) Expression matrices of 73 *Ciona* TF genes during the early (**a**) and late (**b**) gastrula stages. Dark violet indicates expression.

**Supplementary Fig. 3.**
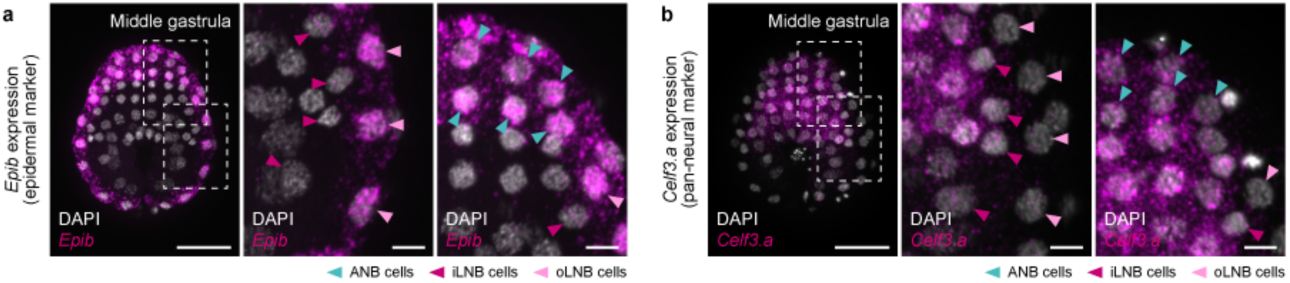
Expression of non-TF markers for epidermis and neural cells in *Ciona* embryos. (**a,b**) *In situ* hybridization of the epidermal marker *Epib* (**a**) and pan-neural marker *Celf3.a* (**b**) in middle gastrula embryos (magenta). Middle and right panels show higher-magnification views. Cells are colored as follows: iLNB (magenta), oLNB (pink), ANB (cyan), neural plate (grey), and pigment-lineage cells (black). Photographs are z-projected image stacks overlaid in pseudocolor. Nuclei are stained with DAPI (grey). Scale bars, 50 μm (left panels), 10 µm (middle and right panels). Brightness and contrast of photographs were linearly adjusted.

**Supplementary Fig. 4.**
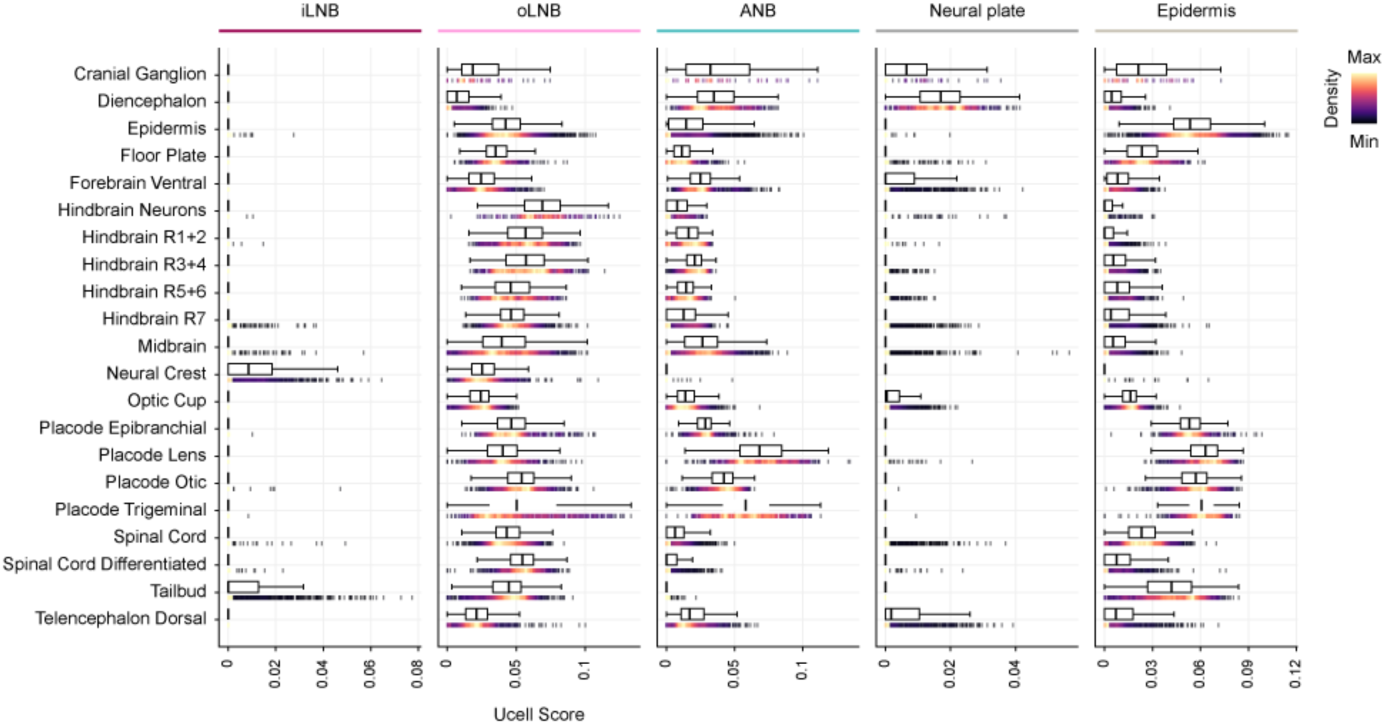
Enrichment of *Ciona* domain gene sets in zebrafish embryos. Rug and box plots visualizing gene set scores for respective domains. Rug plot colors indicate data density. For the box plot, the central line of each box indicates the median, while the lower and upper hinges represent the first (25^th^ percentile) and third (75^th^ percentile) quartiles, respectively. The whiskers extend to the furthest data points that fall within 1.5x the interquartile range from the hinges.

**Supplementary Fig. 5.**
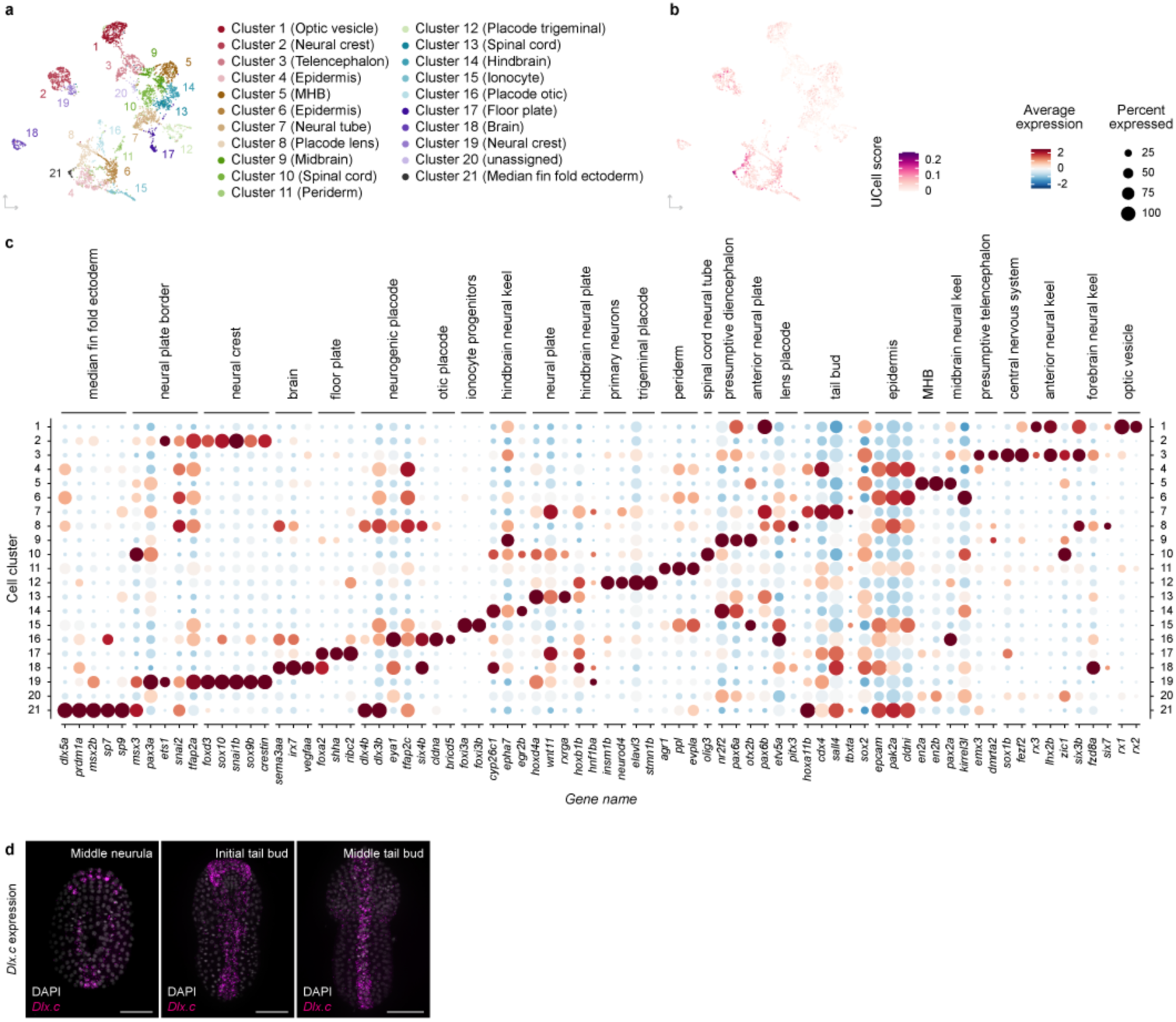
Expression enrichment of orthologs of the *Ciona* oLNB domain-specific gene set in the median fin fold cluster. (**a**) UMAP visualization of single-cell transcriptome data for zebrafish 16-hpf embryos (Lange et al., 2024). Clusters are distinguished by color. (**b**) UCell scores for orthologs of the *Ciona* oLNB domain-specific gene set are mapped onto the zebrafish single-cell transcriptome dataset. (**c**) A dot plot showing expression of marker genes for zebrafish tissues in 16-hpf embryos. Dot sizes and colors indicate percentages of cells that expressed marker genes and average expression, respectively. (**d**) *In situ* hybridization of *Dlx.c*, one of the *Ciona* orthologs for zebrafish *dlx5a*, from the middle neurula to the middle tailbud stage (magenta). Photographs are *z*-projected image stacks overlaid in pseudocolor. Nuclei are stained with DAPI (grey). Scale bars, 50 μm. Brightness and contrast of photographs were linearly adjusted.

**Supplementary Fig. 6.**
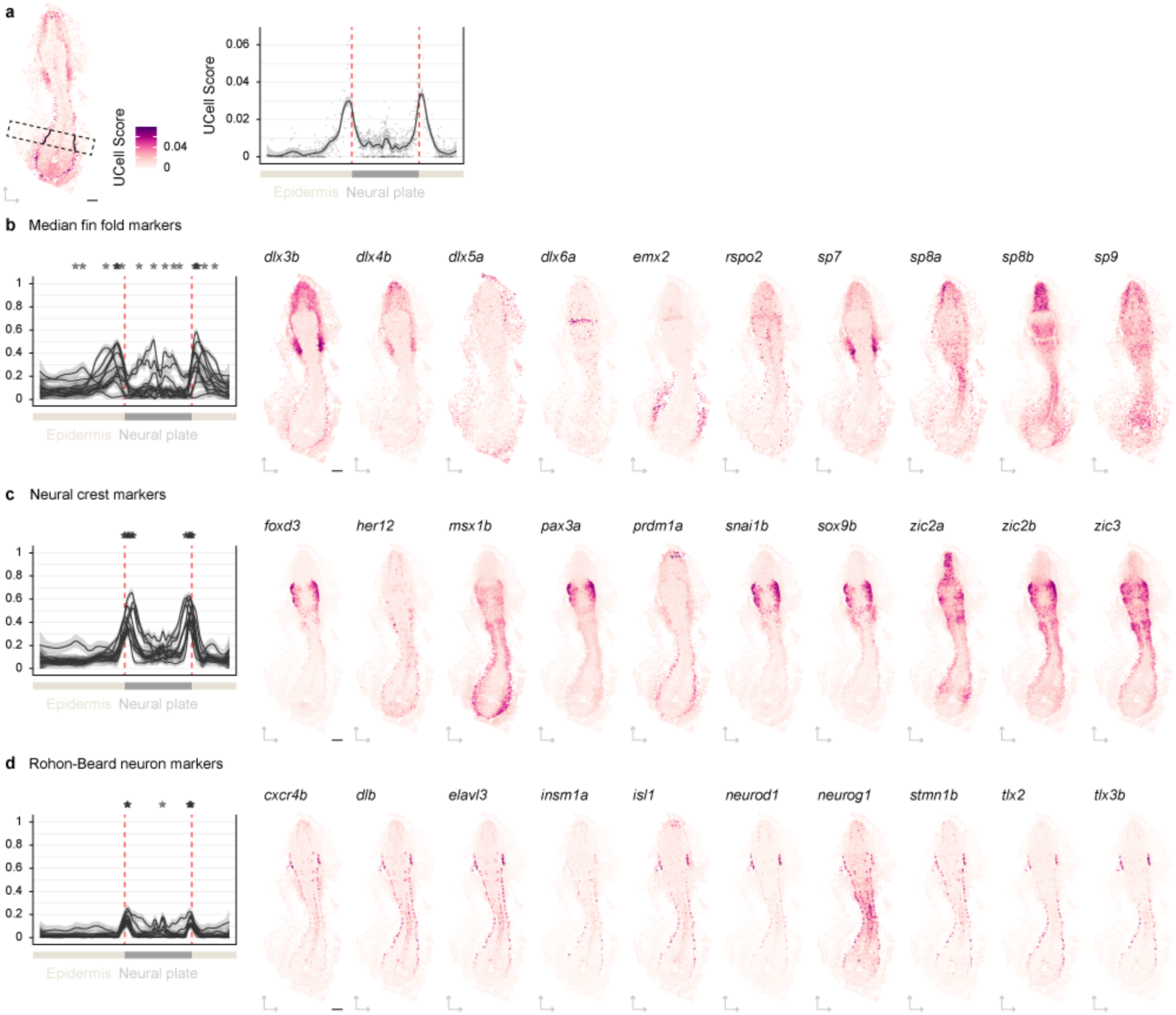
The lateral region of the zebrafish neural plate border can be subdivided into distinct domains. (**a**) Spatial mapping of UCell scores for zebrafish 16-hpf MFF ectoderm-specific gene sets onto the zebrafish spatial transcriptome dataset (left). Black dashed boxes highlight the anterior and posterior regions analyzed in the right panel. Black lines denote boundaries of the neural plate. UCell scores along the normalized mediolateral axis are plotted on the right. Red dashed lines denote the boundaries of the neural plate. (**b,c**) Expression patterns of known markers for neural crest, MFF, and Rohon-Beard neurons in zebrafish spatial transcriptome datasets. (**b-d**) Left: Relative expression of (**b**) MFF ectoderm, (**c**) established neural crest, and (**d**) Rohon-Beard neuron markers along a normalized mediolateral axis in the black dashed box in the **a**. Neural crest, MFF ectoderm, and Rohon-Beard neuron markers are represented in dark red, cyan, and green, respectively. Asterisks denote positions of the peak expression of individual markers on both sides along the normalized mediolateral axis of embryos. Red dashed lines denote the boundaries of the neural plate. Right: Spatial expression patterns of these markers. Scale bar: 100 µm.

